# The activity and functions of soil microbial communities in the Finnish sub-Arctic vary across vegetation types

**DOI:** 10.1101/2021.06.12.448001

**Authors:** Sirja Viitamäki, Igor S. Pessi, Anna-Maria Virkkala, Pekka Niittynen, Julia Kemppinen, Eeva Eronen-Rasimus, Miska Luoto, Jenni Hultman

## Abstract

Increased microbial activity in high-latitude soils due to climate change might lead to higher greenhouse gas (GHG) emissions. However, mechanisms of microbial GHG production and consumption in tundra soils are not thoroughly understood. We analyzed 116 soil metatranscriptomes from 73 sites in the Finnish sub-Arctic to investigate how the diversity and functional potential of bacterial and archaeal communities vary across vegetation types and soil layers. Soils differed in physicochemical conditions, with meadow soils being characterized by higher pH and low soil organic matter (SOM) and carbon/nitrogen ratio whereas dwarf shrub-dominated ecosystems with high SOM and low pH. Actinobacteria, Acidobacteria, Alphaproteobacteria, and Planctomycetes predominated all communities but there were significant differences on genus level between vegetation types, as plant polymer degrading groups were more active in shrub-dominated soils compared to meadows. Given that climate change scenarios predict expansion in dwarf shrubs at high latitudes, our results indicate that the rate of carbon turnover in tundra soils may increase in the future. Additionally, transcripts of methanotrophs were detected in the mineral layer of all soils, potentially moderating methane fluxes from deeper layers. In all, this study provides new insights into possible shifts in tundra microbial diversity and activity with climate change.

## Introduction

The Arctic is one of the regions experiencing the most rapid and severe effects of climate change (IPCC, 2021). Major ecological disturbances have already been observed in Arctic ecosystems and are expected to become more frequent over the coming decades even if anthropogenic greenhouse gas (GHG) emissions are curbed (Post *et al*., 2019). For example, a systematic greening of the Arctic tundra has been observed over the last decades, accompanied by increased plant productivity and the northward and upslope expansion of tall shrubs and trees into this otherwise treeless biome (Frost & Epstein, 2014; Heijmans *et al*., 2022). In addition to regional disturbances, the effects of Arctic climate change might have wider global consequences due to the vast amounts of carbon (C) and nitrogen (N) stored in frozen tundra soils (Mackelprang *et al*., 2011; Johnston *et al*., 2019). Given that warmer temperatures lead to increased rates of soil decomposition and GHG release, Arctic organic matter stocks can contribute to a positive warming feedback loop (Bond-Lamberty *et al*., 2018; Jansson & Hofmockel, 2020).

Microorganisms are important drivers of nutrient cycling in the tundra, and thus the investigation of how microbial communities respond to local environmental variation in tundra soils is essential to predict the impacts of climate change on the GHG budget of this biome (Buckeridge *et al*., 2013; Virkkala *et al*., 2018; Mod *et al*., 2021). Tundra microbial communities are shaped by extreme environmental stressors such as fluctuating temperatures, long periods of sub-zero temperatures, frequent freeze-thaw events, intensive UV radiation, and drought. However, microbial community composition is stable and diverse across seasons in the sub-Arctic tundra, with Acidobacteria being the predominant phylum in acidic soils (Männistö *et al*., 2007, 2013; Pessi *et al*., 2021a). Soil microbes participate in organic matter decomposition, methanogenesis, and methanotrophy in the high-Arctic and permafrost soil ecosystems (e.g., Hultman *et al*., 2015; Schostag *et al*., 2015; Tveit *et al*., 2015), including in peatlands experiencing permafrost thaw (McCalley *et al*., 2014; Singleton *et al*., 2018; Woodcroft *et al*., 2018). However, their functional potential has not been explored in detail due to technical limitations and their vast diversity. This, in turn, hinders a comprehensive understanding of the contribution of tundra microorganisms to and their feedback with climate change. A better understanding of tundra microbial communities and their functions is the key to acquiring process-level knowledge on biogeochemistry and developing accurate models of GHG cycling.

At large geographic scales, the composition of Arctic tundra vegetation is primarily shaped by climate (e.g., mean summer temperature) (Walker *et al*., 2005). However, tundra vegetation is typically heterogeneous at the local level, as growing conditions (microclimate, soil moisture, soil nutrients) vary greatly at small spatial scales (le Roux *et al*., 2013; Kemppinen *et al*., 2021a). Different vegetation types affect soil biotic and abiotic factors which, in turn, influence the local soil microbial community. For example, tundra soils have relatively low pH (4-6) (Hobbie & Gough, 2002; Männistö *et al*., 2007), which is generally one of the most important drivers of microbial community composition and activity (Chu *et al*., 2010). In addition, the quantity and quality of soil organic matter (SOM), N availability, and C/N ratio affect microbial processes and primary production in tundra soils (Koyama *et al*., 2014, Zhang *et al*., 2014). Despite a great local heterogeneity, only fragmentary knowledge exists regarding microbial community composition and activity across different vegetation types in the drier upland tundra, which is a noteworthy ecosystem that covers ca. 90% of the Arctic (Walker *et al*., 2005). This knowledge gap is relevant as most of the Arctic is greening and the typically low-growing Arctic vegetation is being gradually replaced by taller woody plants, a development known as shrubification (Mod & Luoto 2016, Myers-Smith *et al*., 2020; Heijmans *et al*., 2022). Shifts in vegetation and, in particular, shrub expansion across the Arctic tundra are some of the most important ecosystem responses to climate change. These shifts in vegetation potentially alter ecosystem carbon balances by affecting a complex set of soil-plant–atmosphere interactions (Mekonnen *et al*., 2021). In general, decomposition in the Arctic tundra has been slower than plant growth, causing a build-up of detritus in tundra soils. Climate warming may result in carbon loss by accelerating the decomposition of SOM. Future C storage in the Arctic tundra will depend on the balance of C losses from SOM and C storage in plant pools due to higher productivity and changes in plant community assemblages (Weintraub *et al*., 2005).

The warming trend in the Arctic region is alarming but the response of Arctic ecosystems to climate change is only poorly understood. The effects of climate change are particularly complex in tundra ecosystems given their high biotic and abiotic heterogeneity. Thus, elucidating the taxonomic and functional composition of microbial communities across tundra soils is essential to a better understanding of the effects of climate change on the Arctic and potential feedbacks with the global climate system. Here we used metatranscriptomics to investigate the activity of microbial communities during the active growing season across 73 sites in a mountain tundra ecosystem in the Finnish sub-Arctic covering different vegetation types and a wide range of microclimatic and soil nutrient conditions. We aimed at investigating the effect of different vegetation types and soil nutrient conditions on microbial diversity and activity in tundra soils to obtain insights on potential future changes on microbial communities and functions associated with the increased greening of Arctic ecosystems.

## Materials and methods

### Study setting and sampling

The study area is located in Kilpisjärvi, north-western Finland, and extends to the Scandinavian Mountains **(Figure 1)**. The 3 km^2^ area covers parts of two mountains, Mount Saana (69°02′N, 20°51′) and Mount Korkea-Jehkas (69°04′N, 20°79′E) and the valley in between. The elevation range in the study area is 320 m, with the highest point on Mount Saana at 903 m a.s.l. The study area is topographically heterogeneous and part of the sub-Arctic alpine tundra biome. Consequently, the area comprises relatively broad environmental gradients of soil microclimate, moisture, and pH, among others (Kemppinen *et al*., 2021a). The vegetation type is mainly mountain heath dominated by dwarf shrubs such as *Empetrum nigrum* and *Betula nana* and to a lesser extent by *Juniperus communis, Vaccinium vitis-idaea, V. uliginosum*, and *V. myrtillus* (Kemppinen *et al*., 2019). However, due to fine-scale environmental variation and broad gradients, the landscape forms a mosaic of different vegetation types, as both vegetation cover and plant species composition can vary over very short distances (le Roux *et al*., 2013). The soils in the area are mostly poorly developed leptosols with shallow organic layers and occasional podzolization; however, the meadows have soils with thicker organic layers. Permafrost is absent from these soils but can be found in the bedrock above 800 m a.s.l. (King and Seppälä 1987). The average air temperature and precipitation in July for the period 1981–2010 measured at the Kilpisjärvi meteorological station (69°05′N 20°79′E, 480 m a.s.l.) were 11.2°C and 73 mm, respectively (Pirinen et al. 2012).

**Figure 1.**
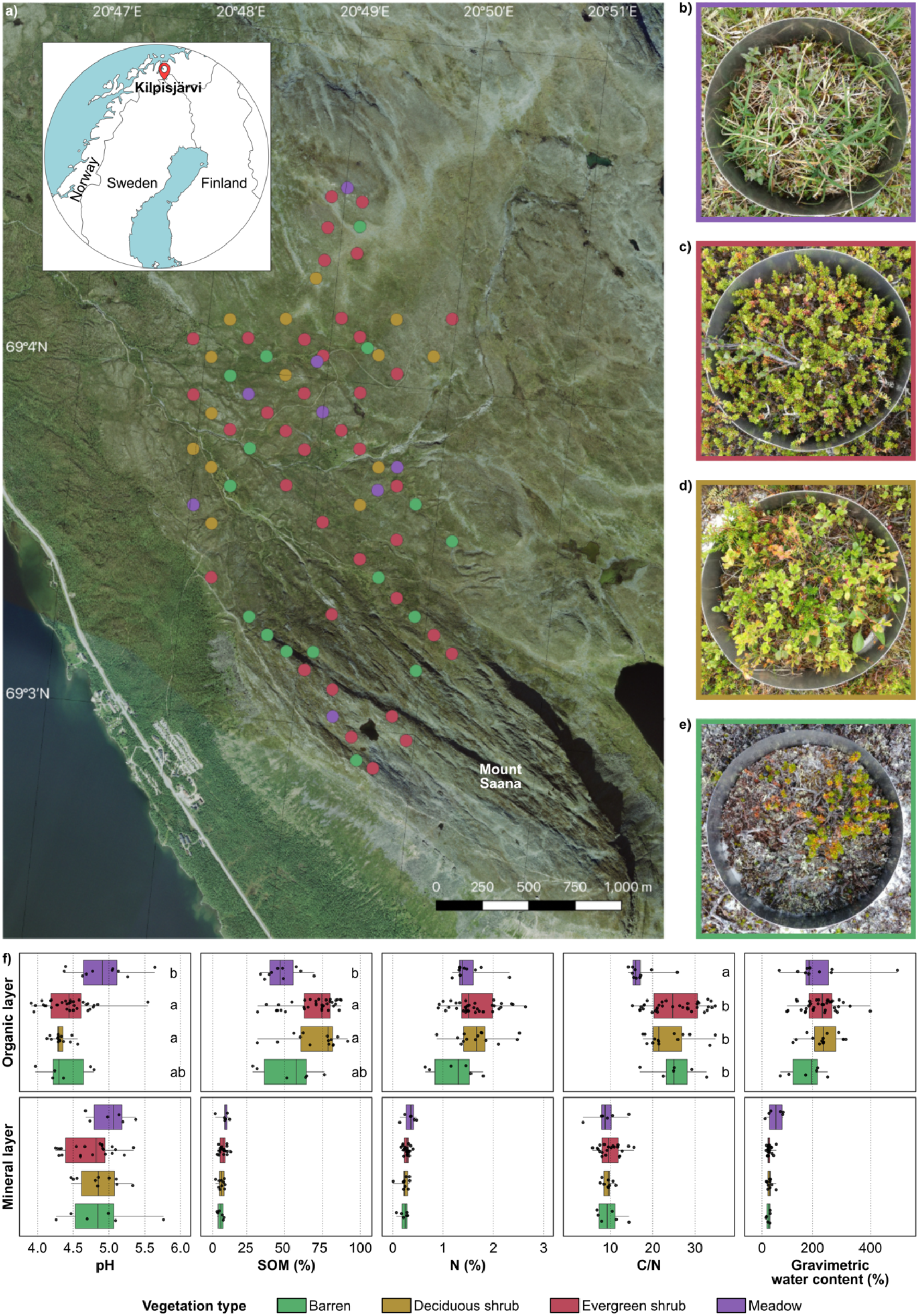
**a)** Map of the sampling sites in Kilpisjärvi, north-western Finland. The inset shows the location of Kilpisjärvi in Fennoscandia. **b–e)** Pictures of the four types of soil vegetation studied (b: meadow; c: evergreen shrub; d: deciduous shrub, e: barren). The number of samples analysed from each vegetation type is indicated. **f)** Soil physicochemical properties across the four vegetation types. Categories with the same letter are not statistically different (one-way ANOVA, p > 0.05). One outlier was removed from the mineral C/N plot.

Samples were collected in July 2017 from 73 sites **(Figure 1a, Supplementary Table 1)**. Sampling sites encompassed four vegetation types, namely barren soil, deciduous shrub, evergreen shrub, and meadow **(Figure 1b-e)**, which were classified according to the Circumpolar Arctic Vegetation Map (Walker *et al*. 2005). All sampling equipment was disinfected with 70% ethanol before and between samples to avoid contamination. The soil was bored with a 50-mm diameter stainless-steel soil corer with a plastic inner casing. When available, both organic and mineral layer sub-samples were collected. Sampling was targeted below the plant roots, with a 5-cm target depth for the organic layer sample, whereas the mineral layer sample was taken from the lowest part of the core from 10-15 cm. The samples were placed in a Whirl-Pak sampling bag (Nasco, Fort Atkinson, WI, USA) with a metal spoon and immediately frozen on dry ice and kept frozen at –80°C until nucleic acid extraction. Samples were collected from 73 sites, from which 116 metatranscriptomes were sequenced.

### Soil physicochemical data

For the analysis of soil physicochemical properties, approximately 0.2 dm^3^ of soil was collected with a steel cylinder and stored at 4 °C. The soils were lyophilized according to the Finnish standard SFS300 and pH was analysed according to international standard ISO10390. SOM content was determined by loss on ignition analysis according to the Finnish standard SFS3008. CNS analysis (carbon, nitrogen, and sulphur) was performed with a Vario Micro Cube analyser (Elementar, Langenselbold, Germany). For this, mineral samples were sieved through a 2-mm plastic sieve and the organic samples were homogenized by hammering the material into smaller pieces. Differences in soil physicochemical properties between vegetation types were assessed using the Kruskal-Wallis test followed by the pairwise Wilcoxon test with Bonferroni correction (functions *kruskal.test* and *pairwise.wilcox.test*, R-core package).

### Nucleic acid extraction

Three replicate nucleic acid extractions were performed for each sample. Samples were kept on ice during weighing and extraction and all steps were performed promptly with nuclease-free labware to avoid RNA degradation. Solutions and water were treated with 0.1% diethylpyrocarbonate. Nucleic acids were extracted using a modified hexadecyltrimethylammonium bromide (CTAB), phenol-chloroform, and bead-beating protocol (Griffiths *et al*., 2000; DeAngelis *et al*., 2009). On dry ice, approximately 0.5 g of frozen soil was transferred to a 2-mL Lysing Matrix E tube (MP Biomedicals, Heidelberg, Germany) and 0.5 mL of CTAB buffer (consisting of equal amounts of 10% CTAB in 1 M sodium chloride and 0.5 M phosphate buffer in 1 M NaCl), 50 μl 0.1 M ammonium aluminium sulfate (NH_4_(SO_4_)_2_ ·2 H_2_O), and 0.5 mL phenol:chloroform:isoamyl alcohol (25:24:1) was added. After bead-beating with FastPrep (MP Biomedicals, Heidelberg, Germany) at 5.5 m s^−1^ for 30 s, 0.5 mL chloroform was added. Nucleic acids were precipitated with polyethylene glycol 6000 (PEG6000, 30% in 1.6 M NaCl) and washed with ethanol. Nucleic acids were extracted again from the leftover soil pellet to maximize yields. Nucleic acids were resuspended in 25 μl of Buffer EB and 250 μl Buffer RLT with β-mercaptoethanol added. Buffers EB and RLT were from an AllPrep DNA/RNA Mini Kit (Qiagen, Hilden, Germany). All centrifugations were performed at 4°C. Finally, RNA and DNA were purified with AllPrep DNA/RNA Mini Kit (Qiagen, Hilden, Germany), where RNA was treated with DNAse I. The amount and integrity of RNA were assessed on a Bioanalyzer RNA 2100 Nano/Pico Chip with Total RNA Assay (Agilent, Santa Clara, CA, USA). To ensure that RNA was DNA-free, a check-up PCR with universal primers and gel electrophoresis was performed. Triplicates were pooled by combining equal amounts of RNA from each triplicate. Ribosomal RNA was not depleted and thus the total RNA approach was used (Urich *et al*., 2008).

### Sequencing

Complementary DNA (cDNA) libraries were constructed with the Ultra II RNA Library Prep Kit for Illumina (New England Biolabs, Ipswich, MA, USA). cDNA concentrations were measured using a Qubit fluorometer with a dsDNA BR/HS kit (Invitrogen, Carlsbad, CA, USA). Before sequencing, the libraries were analysed with Fragment analyzer (Advanced Analytical, Ames, IA, USA) and small cDNA fragments were removed to avoid primer binding to the flow cell and to reduce cluster density. Single-end sequencing was performed on an Illumina NextSeq 500 (Illumina, San Diego, CA, USA) with 150 cycles at the Institute of Biotechnology, University of Helsinki, Finland.

### Metatranscriptomic data processing and analysis

One sample (site 11202, organic layer) yielded a much higher than average amount of reads (80.4 million) and was randomly reduced to 4 million reads with seqtk v. 1.3 (https://github.com/lh3/seqtk). The quality of the sequences was assessed with FastQC v. 0.11.5 (https://www.bioinformatics.babraham.ac.uk/projects/fastqc) and MultiQC v. 1.3 (Ewels *et al*., 2016). Trimming and quality filtering was performed with Cutadapt v. 1.10 (Martin, 2011) applying a quality cut-off of 25 and a minimum adapter overlap of 10 bp. Metaxa2 v. 2.1.3 (Bengtsson-Palme *et al*., 2015) was used to identify reads mapping to the small subunit (SSU) rRNA. These were then classified against the SILVA database release 132 (Quast *et al*., 2013) using the mothur v. 1.40.5 classify.seqs command with a confidence cut-off of 60% (Schloss *et al*., 2009). The taxonomy of abundant taxa was manually updated according to the Genome Taxonomy database (Parks *et al*., 2018, 2020). For the analysis of protein-coding genes, reads were mapped to the Kyoto Encyclopedia of Genes and Genomes (KEGG) Prokaryote database release 86 (Kanehisa & Goto, 2000) using DIAMOND blastx v. 2.1.3 (Buchfink *et al*., 2015) with an E-value cut-off of 0.001. The KEGG orthology (KO) identifier of the best hit was assigned to each read and mapped to the KEGG module hierarchy, and spurious pathways were removed using MinPath v. 1.4 (Ye & Doak, 2009). Since genes for methane and ammonia oxidation (*pmo* and *amo*, respectively) are not distinguished in KEGG we used blastx (Altschul *et al*., 1990) to compare putative *pmoA-amoA* genes against a manually curated database of five PmoA and nine AmoA sequences from both bacteria and archaea **(Supplementary Figure 1)**.

Statistical analyses and visualization were performed with R v. 3.6.2 (R Core Team 2020). For multivariate analyses, taxonomic (genera abundances) and functional (KO abundances) matrices were transformed into Bray-Curtis distance matrices (function *vegdist*, R-package vegan v. 2.5-6; https://github.com/vegandevs/vegan). Community-wide differences between vegetation types and soil layers were tested with permutational analysis of variance (PERMANOVA) (function *adonis*, R-package vegan v. 2.5-6) followed by pairwise PERMANOVA (function pairwise.perm.manova, R-package RVAideMemoire v.0.9-78; https://cran.r-project.org/web/packages/RVAideMemoire). Differences in community structure were visualized using principal coordinates analysis (PCoA) (function *ordinate*, R-package Phyloseq v. 1.30.0; McMurdie & Holmes, 2013). The relationship between community structure and soil physicochemical properties was assessed using distance-based redundancy analysis (db-RDA) with forward selection (functions *capscale* and *ordistep*, R-package vegan v. 2.5-6). Soil physicochemical data was log-transformed prior to the analysis. Differences in the abundance of individual bacterial and archaeal genera and functional genes between vegetation types were tested with one-way analysis of variance (ANOVA) (functions *lm* and *aov*, R-core package) followed by pairwise t-test (function *pairwise.t.test*, R-core package).

### Data availability

Sequences have been deposited in the European Nucleotide Archive under accession number PRJEB45463.

## Results

### Soil properties across vegetation types

We observed a high variability in soil properties between samples from the organic and mineral layers and across vegetation types. SOM content varied from 2% to 92%, gravimetric soil water concentration from 9% to 432%, and pH from 3.9 to 5.8 **(Figure 1f)**. pH was generally higher in the mineral layer whereas SOM, N, C/N ratio, and gravimetric water content were higher in the organic layer. Soil properties also differed between vegetation types (Kruskal-Wallis test; *P* < 0.01). In the organic layer, meadow sites were less acidic and contained less SOM than deciduous or evergreen shrubs. Meadows also had a lower C/N ratio than shrubs or barren soils. The physicochemical properties of the mineral layer did not differ significantly between vegetation types. Varying degrees of collinearity between the physicochemical variables was observed **(Supplementary Figure 2)**.

### Microbial community composition along the tundra landscape

We obtained 281.1 million sequence reads using a total RNA metatranscriptomic approach (Urich *et al*., 2008). First, we analyzed reads corresponding to the small subunit ribosomal RNA (SSU rRNA). SSU rRNA represented 35 ± 2% (mean ± standard deviation) of the reads. Over 80% of the SSU rRNA sequences were bacterial, 0.1 % archaeal (mostly Thaumarchaea), and 19% were of eukaryotic origin, with fungal SSU rRNA representing 12% of the sequences (Ascomycota, 8%; Basidiomycota, 3%). Furthermore, 0.2% of the sequences were recognized as SSU rRNA but could not be assigned unambiguously to a specific domain.

The predominant bacterial groups were assigned to the phylum Actinobacteria (27 ± 9% of the sequences; orders Acidothermales and Solirubrobacterales), phylum Acidobacteria (17 ± 3%; orders Acidobacteriales and Solibacterales), class Alphaproteobacteria (16 ± 3%; orders Rhizobiales and Acetobacterales), and phylum Planctomycetes (14 ± 3%; order Gemmatales) **(Figure 2)**. Classes Deltaproteobacteria and Gammaproteobacteria, as well as phyla Chloroflexi and Verrucomicrobia, were also abundant. The most abundant genera were *Acidothermus* (phylum Actinobacteria; 13 ± 5.7%) and *Ca*. Solibacter (phylum Acidobacteria; 3.11 ± 0.83%), followed by *Bryobacter* (phylum Acidobacteria; 2.06 ± 0.38%), *Pajaroellobacter* (class Deltaproteobacteria; 1.64 ± 0.55%), and *Roseiarcus* (class Alphaproteobacteria; 1.48 ± 1.07%) **(Figure 3, Supplementary Table 2)**.

**Figure 2.**
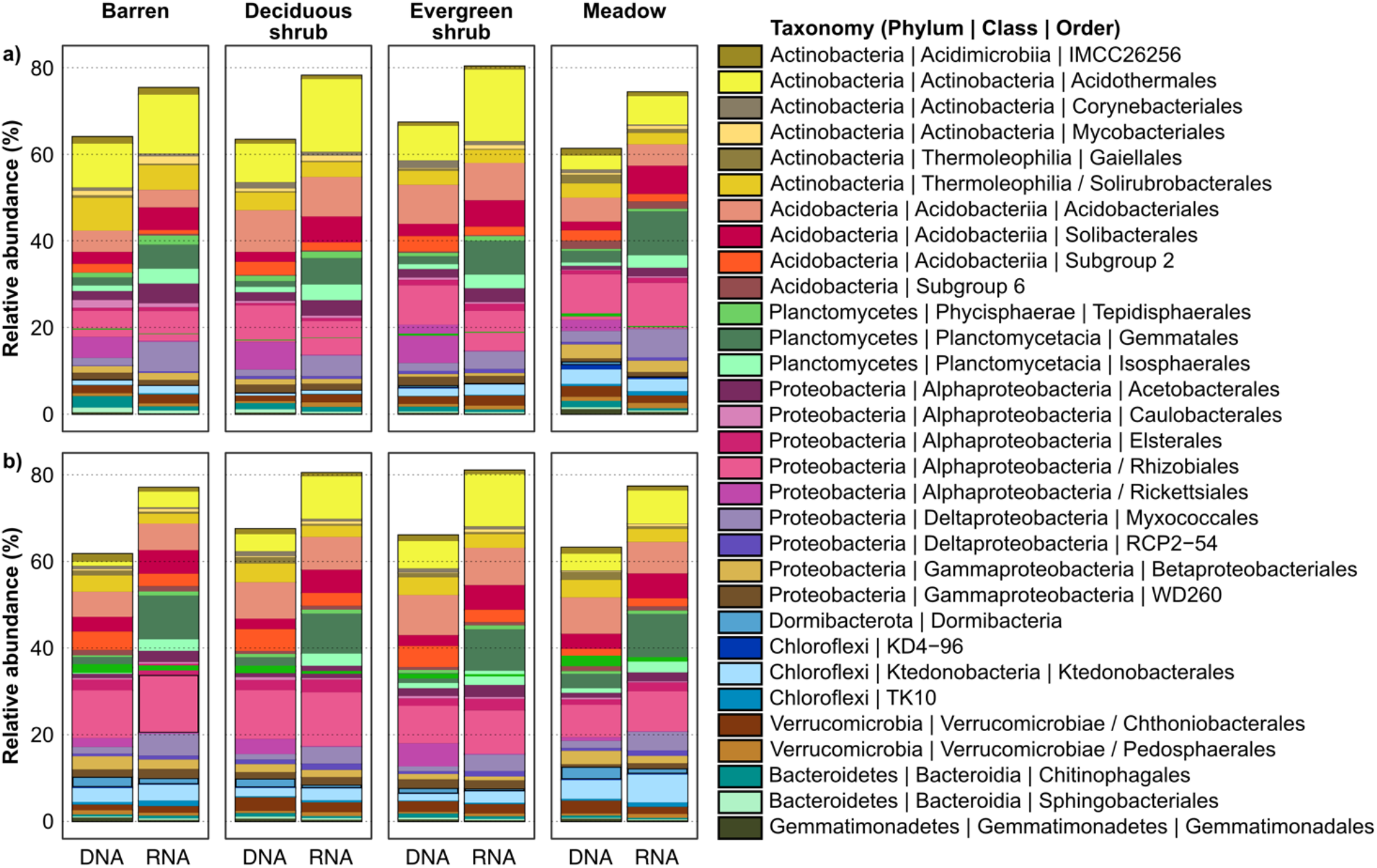
Relative abundances of bacterial orders in metatranscriptomes (RNA) and metagenomes (DNA) of samples from the **a)** organic and **b)** mineral layer. Samples from the same vegetation type were pooled and unclassified taxa were removed.

**Figure 3.**
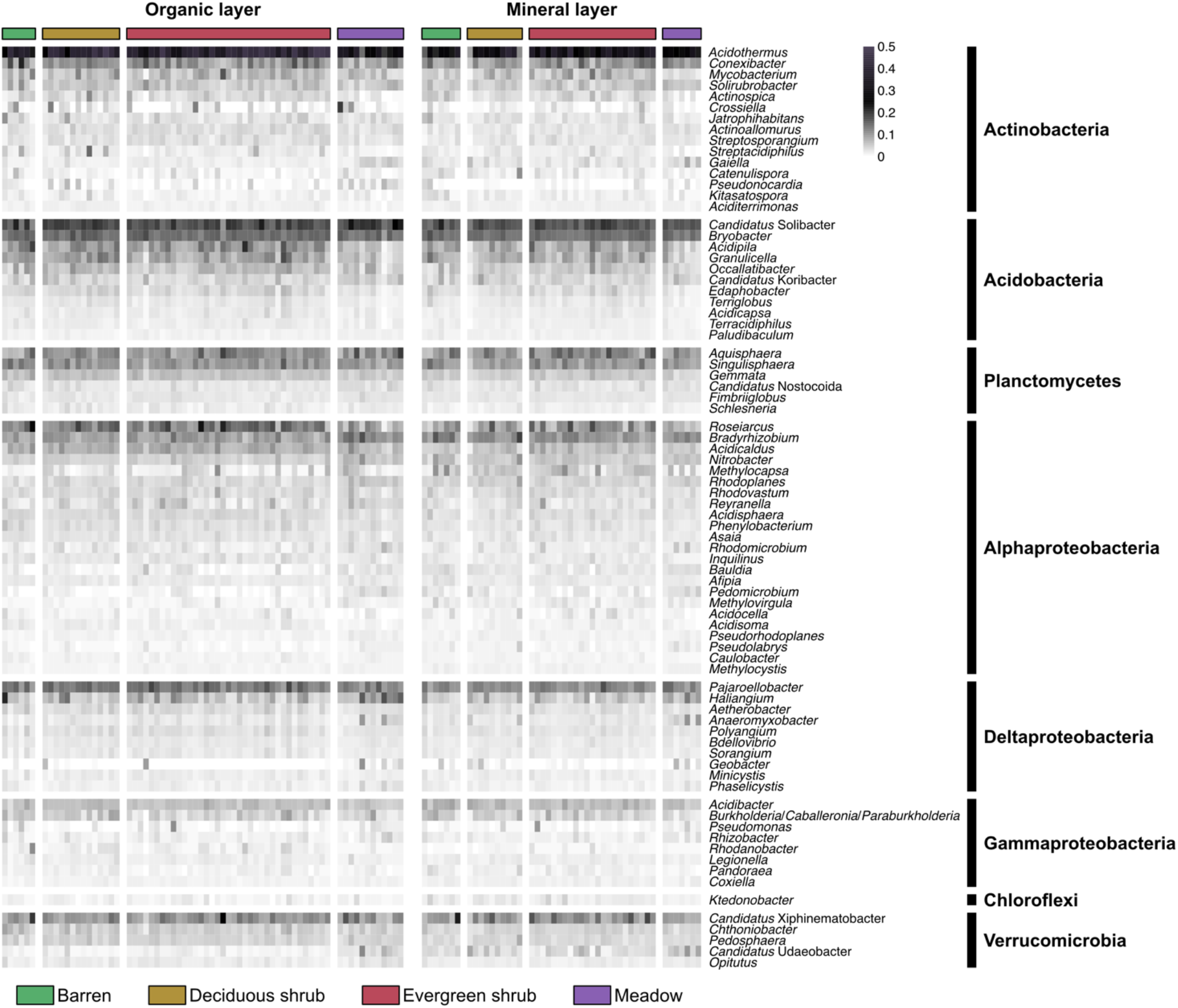
Relative abundances of the 80 most abundant bacterial genera across the samples. Abundances were square root-transformed to improve visualization.

When comparing SSU rRNA sequences from metatranscriptomes (i.e. RNA, representing active microbes) to sequences from metagenomes (i.e. DNA, representing the whole microbial community) (Pessi *et al*., 2021a), the most abundant microbial groups were largely the same but with notable differences **(Figure 2, Supplementary Figure 3a)**. Orders with greater relative abundance in the metatranscriptomes included Gemmatales (Planctomycetes), Acidothermales (Actinobacteria), Solibacterales (Acidobacteria), and Myxococcales (Deltaproteobacteria), whereas Subgroup 2 (Acidobacteria), Rickettsiales (Alphaproteobacteria), Dormibacterota, Chitinophagales (Verrucomicrobia), and Gemmatimonadales (Gemmatimonadetes) were more abundant in the metagenomes. In the metatranscriptomes, the orders with more than 1% relative abundance accounted together for 78% of the communities compared to 64% in the metagenomes. Additionally, more than 1500 bacterial and archaeal genera were identified from the metatranscriptomes and less than 500 bacterial and archaeal genera from the metagenomes.

### Shifts in microbial community composition across the different vegetation types

Genus-level community structure was significantly different in the organic and mineral layers (PERMANOVA; *R^2^* = 0.07; *P* < 0.001) **(Supplementary Figure 3b)**. Interestingly, communities also differed significantly across the four different vegetation types both in the organic (*R^2^* = 0.16; *P* < 0.001) and mineral layers (*R^2^* = 0.13; *P* < 0.01). Pairwise analyses revealed that the communities in the organic layer of meadow sites were significantly different from all other vegetation types, whereas communities from the mineral layer differed only between the meadow and evergreen shrub sites **(Supplementary Table 3)**.

Genus-level comparisons were conducted for the metatranscriptomes across vegetation types and soil layers. For this, we considered only abundant genera (i.e. genera with a mean abundance at least twice the mean of all genera). In the organic layer, the alphaproteobacterial genera *Bradyrhizobium, Nitrobacter, Rhodoplanes*, and *Rhodomicrobium* were significantly more abundant in samples from the meadow sites than sites with other vegetation types (ANOVA; *P* < 0.01) **(Figure 4a)**. The same was observed for other taxa, namely *Gaiella* (Actinobacteria), *Ca*. Nostocoida (Planctomycetes), “HSB_OF53-F07” (Chloroflexi), *Anaeromyxobacter* (Deltaproteobacteria), *Gemmatimonas* (Gemmatimonadetes), “R4B1” (Acidobacteria), *Ca*. Udaeobacter and “ADurb.Bin063-1” (Verrucomicrobia). On the other hand, *Acidothermus* (Actinobacteria) was less abundant in the meadow than the shrub sites, whereas *Roseiarcus* (Alphaproteobacteria), the acidobacterial genera *Acidipila, Granulicella*, and *Edaphobacter*, as well as *Gemmata* (Planctomycetes) were less abundant in the meadow sites compared to all other vegetation types. In the mineral layer, “HSB_OF53-F07” (Chloroflexi), *Anaeromyxobacter* (Deltaproteobacteria), “Adurb.Bin063-1” (Verrucomicrobia), and “R4B1” (Acidobacteria) were more abundant in the meadows compared to all other vegetation types, whereas *Bauldia* (Alphaproteobacteria) was more abundant in the meadows only in relation to the shrub sites **(Figure 4b)**.

**Figure 4.**
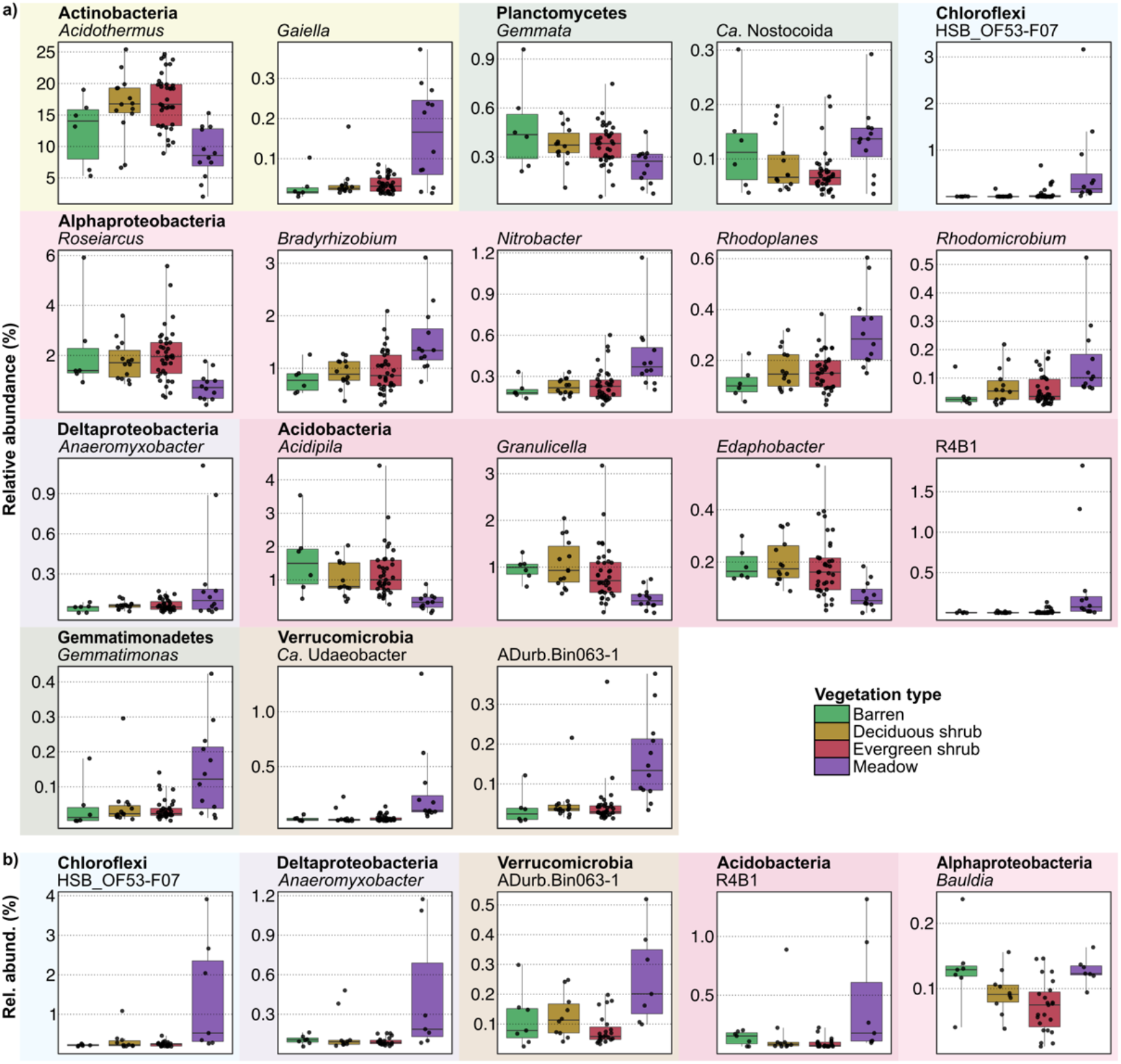
Boxplots showing abundant genera (mean abundance larger than the twofold mean of all genera) that were differentially active across vegetation types in **a)** organic layer and **b)** mineral layer (one-way ANOVA, *P* < 0.01).

We used db-RDA with forward selection to investigate which factors underly the observed differences in microbial community structure. In the mineral layer, the best model included vegetation, pH, and gravimetric water content (*R^2^* = 0.25; *P* < 0.05), whereas in the organic layer the best model included vegetation, pH, and C/N ratio (*R^2^* = 0.28; *P* < 0.05). However, it is important to note that there are varying degrees of collinearity between the variables selected by the forward selection procedure and other variables **(Supplementary Figure 2)**. For example, gravimetric water content in the mineral layer and pH and C/N ratio in the organic layer were correlated with SOM, C, and N content (–0.5 ≤ *r* ≥ 0.5). Thus, in these cases, the variables selected by the model should be considered, to some extent, as a proxy of the intercorrelated variables.

### Microbial community functions across vegetation types

Differences in protein-coding gene composition between samples from the organic and mineral layers were small (PERMANOVA; *R^2^* = 0.03; *P* < 0.001). Community structure based on protein-coding genes was also significantly different between vegetation types in the organic layer (*R^2^* = 0.10; *P* < 0.001), with communities from the meadow sites differing from shrub and barren sites and evergreen shrub communities differing from barren sites. No significant differences were observed between vegetation types in the mineral layer.

Genes with a KEGG classification represented only a small fraction (1.39 ± 0.27%) of the protein-encoding genes. While genes with no KEGG classification were not analysed in this study, they corresponded mostly to genes encoding proteins with unknown function **(Supplementary Figure 4)**. The most abundantly transcribed genes that were mapped to KEGG pathways are involved in genetic information processing, including i) folding, sorting, and degradation, ii) transcription, and iii) metabolism **(Supplementary Figure 5)**. ABC transporter genes were widely transcribed, including genes encoding transport system substrate-binding proteins for ribose (*rbsB*), D-xylose (*xylF*), and sorbitol/mannitol (*smoE, mtlE*), and multiple sugar transport system ATP-binding proteins (*msmX, msmK, malK, sugC, ggtA, msiK*), indicating the decomposition of plant polymers. Other widely transcribed ABC transporters included genes for branched-chain amino acid transport system proteins (*livKGFHM*) and urea transport system substrate-binding proteins (*urtA*). In addition to transporters, the chaperone genes *groeL* and *dnaK* and the cold shock protein gene *cspA* involved in survival in cold temperatures were among the most transcribed genes across all samples. The gene *coxL/cutL* encoding the large subunit of the aerobic carbon-monoxide dehydrogenase enzyme was also widely expressed, as well as two genes involved in nitrogen uptake, namely glutamine synthetase (*glnA*) and ammonium transporter (*amt*).

Given the high abundance of genes involved in the transport of carbohydrates, we expanded our analysis to other genes related to C cycling and metabolism **(Supplementary Figure 6)**. Interestingly, the *pmoABC-amoABC* genes involved in methane/ammonia oxidation were significantly more transcribed in the mineral layer than in the organic layer (ANOVA; *pmoA: R^2^* = 0.13, *pmoB: R^2^* = 0.09, *pmoC: R^2^* = 0.12; *P* < 0.001) **(Figure 5)**. To further distinguish between the closely related *amoA* and *pmoA* genes, we performed a blastx analysis against a manually curated database of PmoA and AmoA sequences from different organisms. This indicated that 96% of the sequences identified as *pmoA*-*amoA* using the KEGG database correspond to the *pmoA* gene, indicating methane oxidation by the particulate methane monooxygenase (pMMO). However, the soluble methane monooxygenase (sMMO) genes *mmoXYZBCD*, as well as the genes *mxaF* or *xoxF* which encode methanol dehydrogenases for methanol oxidation to formaldehyde, were in general not transcribed **(Figure 5)**. Genes for formaldehyde assimilation using the serine pathway, including *glyA* encoding the enzyme glycine hydroxymethyltransferase, were transcribed. This indicates the utilisation of the serine cycle instead of the ribulose monophosphate (*RuMP*) cycle for formaldehyde assimilation in these microbial communities.

**Figure 5.**
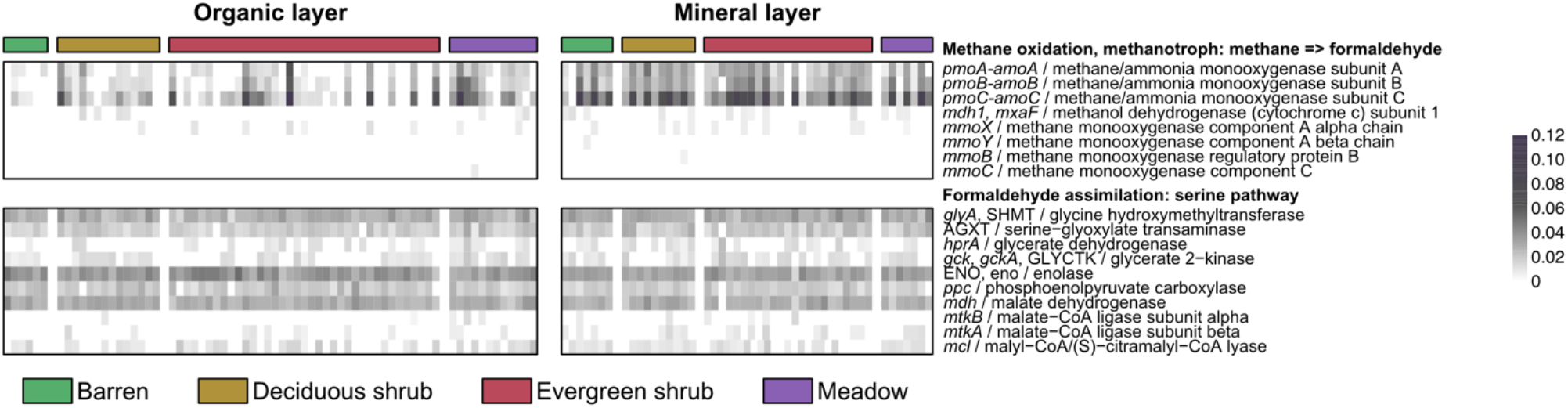
Relative abundances of genes involved in methane oxidation and the serine pathway of formaldehyde assimilation. Abundances were square root-transformed to improve visualization.

**Figure 6.**
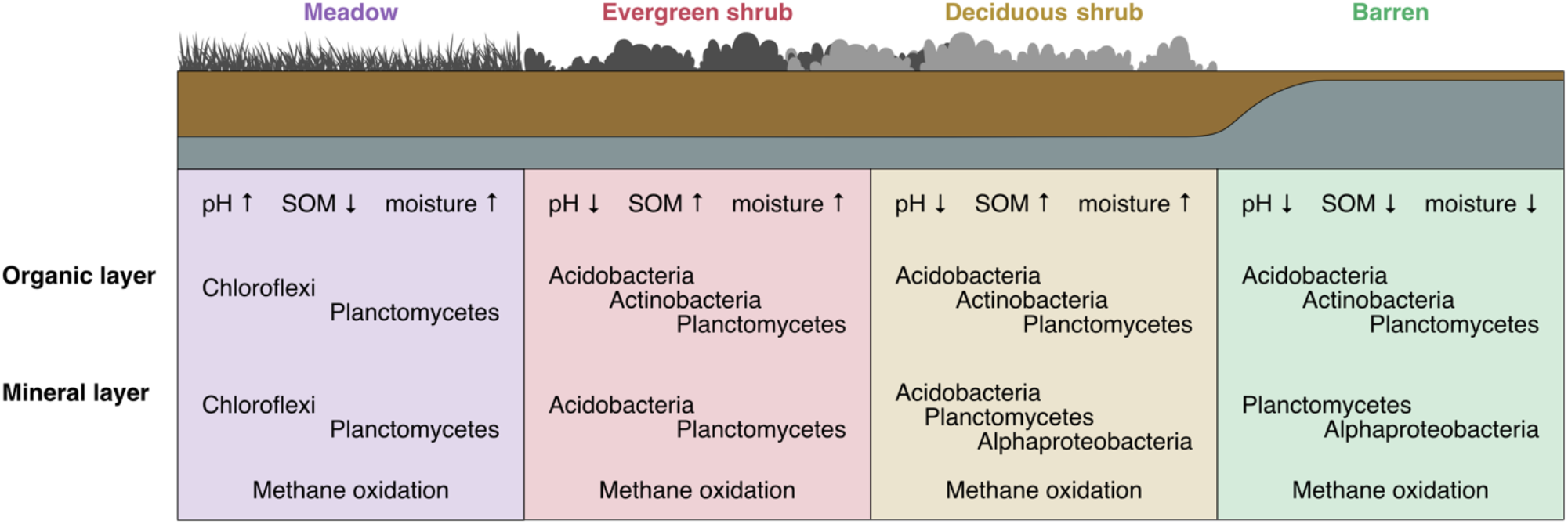
A conceptual figure on the implications of the study. The arrows denote the change in the measured environmental variables and the microbial phyla active in the sites are marked to each vegetation type and soil layer.

## Discussion

We analysed over 100 soil metatranscriptomes across a sub-Arctic tundra landscape to investigate how microbial community composition and their functions vary across soil layers and vegetation types. Soil physicochemical composition varied according to vegetation type in the organic layer, but not in the mineral layer. This is likely related to vegetation being the primary source of material for the organic layer, whereas the mineral layer properties are more affected by bedrock and soil texture, among other factors (Jenny 1941; Haichar *et al*., 2008). Thus, environmental conditions are more homogeneous in the mineral layer, leading to more uniform microbial communities irrespective of vegetation cover as observed in the present study. Differences in soil properties between vegetation types were more pronounced in the organic layer, with the meadows differing significantly from the other vegetation types by higher pH and lower SOM and C/N ratio. As revealed by our multivariate analyses, these factors were significantly associated with differences in community structure observed between vegetation types, which can be presumably linked to differences in SOM quality. Evergreen shrubs such as *Empetrum nigrum*, which is the dominant plant species in the study area, produce recalcitrant, acidic, and slowly decomposing litter. On the other hand, meadows are dominated by forbs, grasses, and sedges, which produce litter that decomposes faster and has higher nutrient concentrations with lower C/N ratio (Hobbie *et al*., 2000; Eskelinen *et al*., 2009).

Actinobacteria, Acidobacteria, Alphaproteobacteria, and Planctomycetes were the most active phyla in all vegetation types which is consistent with previous studies from Arctic regions (Männistö *et al*., 2007; Hultman *et al*., 2015; Taş *et al*., 2018; Tripathi *et al*., 2019, Ivanova *et al*., 2018). These phyla, excluding Planctomycetes, were also among the most abundant in the metagenomics dataset from these samples (Pessi *et al*., 2021a). Archaea, which represented only 0.1% of the transcripts, consisted mostly of Thaumarchaea in the mineral layer as previously observed (Lu *et al*., 2017; Shao *et al*., 2019; Pessi *et al*., 2021b). Communities in both the organic and mineral layers and across all vegetation types were dominated by aerobic acidophilic genera that play a role in the degradation of plant organic matter, including *Acidothermus, Ca*. Solibacter, and *Bryobacter* (Mohagheghi *et al*., 1986, Ward *et al*., 2009, Kulichevskaya et al., 2010). *Acidothermus* was the most abundant active genus overall and was significantly more abundant in the shrub sites. *Acidothermus cellulolyticus*, the only described species in this genus, is a thermophilic, acidophilic, and cellulolytic species first isolated from an acidic hot spring (Mohagheghi *et al*., 1986). Furthermore, these microorganisms tolerate temperature and moisture fluctuation and low-nutrient conditions, which are characteristic of tundra soils (Ward *et al*., 2009; Rawat *et al*., 2012). Interestingly, genome analysis of *Ca*. Solibacter revealed not only the ability to utilise complex plant cell-wall polysaccharides and simple sugars but also carbon monoxide (CO), a toxic gas, as a complementary energy source in a mixotrophic lifestyle (Ward *et al*., 2009). Indeed, the *coxL/cutL* gene encoding the carbon monoxide dehydrogenase enzyme responsible for the oxidation of CO was widely expressed in the present study. In addition, *Ca*. Solibacter and *Bryobacter* are facultative anaerobes that play a role in the nitrogen cycle as they harbour candidate nitrite and nitric oxide reductases (Pessi *et al*., 2021a). In general, our results show that the dominant active microorganisms in the tundra soils studied here are versatile degraders of plant polymers with the ability to thrive in fluctuating conditions and have potential roles in the C and N cycle.

Our results evidenced a link between soil microbial community composition/activity and vegetation via physicochemical factors such as pH and SOM content. Meadow soils, which were characterized by higher pH and lower SOM content and C/N ratio harboured distinct microbial communities compared to the other vegetation types. Most of the genera that were abundant in the meadows are poorly known, and more research is required to understand their roles in this ecosystem. Interestingly, members of *Gaiella, Bradyrhizobium, Ca*. Udaeobacter, and *Gemmatimonas* have been implicated in the cycling of the atmospheric gases H_2_, CO_2_, and N_2_O (Lepo *et al*., 1980; Park *et al*., 2017; Severino *et al*., 2019, Willms *et al*., 2020). Shrub soils, characterized by lower pH and higher SOM content, had a higher abundance of the Acidobacteria genera *Acidipila, Granulicella*, and *Edaphobacter*. These genera most likely have a role in degrading plant-derived organic matter in these shrub soils, as seen in other acidic upland soils (Pankratov & Dedysh, 2010; Männistö *et al.;* 2013, Ivanova *et al*., 2020, 2021). Altogether, our results indicate that shrub soils have a higher abundance of chemoorganotrophs that degrade complex plant polymers, whereas meadows harbour also microbial groups that are not solely dependent on plant-derived organic matter.

Interestingly, our results evidenced a potential for methane oxidation in the mineral layer, as the *pmo* gene and genes for the serine pathway were transcribed together with the activity of methanotrophic bacteria such as *Methylocapsa*. Of the known *Methylocapsa* species, *M. gorgona* grows only on methane, whereas *M. acidiphila* and *M. palsarum* grow also on low methanol concentrations and *M. aurea* on methanol and acetate (Dedysh *et al*., 2002, 2015; Dunfield *et al*., 2010; Tveit *et al*., 2019). A recent *in situ* ^13^CH_4_-DNA-SIP enrichment study showed that *Methylocapsa* were the dominant active methane oxidizers in high Arctic soil (Altshuler *et al*., 2022). Based on the results shown here and by others (e.g., Belova *et al*., 2020), *Methylocapsa* could have a significant role as methane oxidizers in sub-Arctic tundra soils. Moreover, our results showed similar activity levels of methane oxidizers in the mineral layer across all vegetation types, which suggests that methane oxidation is not dependent on vegetation cover in deeper soil layers. Future studies employing e.g. stable isotope probing (SIP) could shed light on the regulation of methane oxidation in tundra soils.

In this study, we investigated the microbial activity across different vegetation types (barren, deciduous shrub, evergreen shrub, and meadow) in tundra soils to understand how future changes in vegetation cover such as shrubification may affect the microbial community diversity and activity. In the dwarf shrub-dominated tundra, shrubs influence microclimate and soil moisture, which can lead to cooling and drying of the soils in the growing season (Kemppinen *et al*., 2021b). Our findings indicate that plant polymer-degrading microorganisms would be active in these conditions. However, the overall greening of the Arctic is more complex, as graminoids instead of shrubs are increasing in colder parts of the region (Elmendorf *et al*., 2012). In addition, with increasing temperatures, high latitudes will receive more precipitation as rainfall across the Arctic (Bintanja & Andry 2017), which will likely affect microbial communities which are strongly reliant on soil moisture resources (Evans *et al*., 2022). Therefore, the consequences of macroclimatic changes on soil moisture are not straightforward. Yet, it is evident that, in addition to the direct effects on microbial communities, future moisture conditions will have a strong effect also in shrubification (Ackerman *et al*., 2016), plant diversity, and assemblages (le Roux *et al*., 2013), and overall ecosystem functions (Bjorkman *et al*., 2018) in the tundra, including microbial mediated processes. In all, despite the overall similarity in bacterial community composition in this study, shrubs had a more abundant and active community of potential degraders of plant-derived organic matter. Therefore, we hypothesize that if shrub soils become more prevalent, heterotrophic microbial activity may increase and lead to increased CO_2_ fluxes.

## Acknowledgments

The study was funded by the Academy of Finland (Grant 1314114) and the University of Helsinki’s three-year grant to JH. SV was funded by the Doctorate Program in Microbiology and Biotechnology. PN was funded by Kone Foundation and Nessling Foundation. JK was funded by the Arctic Interactions at the University of Oulu and Academy of Finland (grant no. 318930, Profi 4). AMV acknowledges Gordon and Betty Moore Foundation (Grant 8414). Permission to perform fieldwork was granted by Metsähallitus. We acknowledge the CSC – IT Center for Science for the computational resources and Mr. Kimmo Mattila for assistance. We acknowledge the Kilpisjärvi Biological Station and staff and members of the BioGeoClimate Modelling Lab for use of the premises and assistance with fieldwork. We would like to thank the anonymous reviewers for their valuable contribution to improving the manuscript.

---

**S1 Supplementary Table S1.** Sampling sites, their coordinates and soil type sampled from (o: organic, m: mineral).

| site | point_x | point_y | org_min |
| --- | --- | --- | --- |
| 19 | 492001.7163 | 7661066.296 | om |
| 25 | 491972.6377 | 7661365.536 | om |
| 31 | 491943.56 | 7661661.864 | om |
| 37 | 491915.594 | 7661960.17 | om |
| 91 | 492137.6413 | 7660682.24 | om |
| 97 | 492108.5998 | 7660978.766 | om |
| 103 | 492081.1879 | 7661276.182 | om |
| 109 | 492050.6532 | 7661570.414 | om |
| 115 | 492021.724 | 7661871.922 | om |
| 181 | 492189.9563 | 7661189.969 | o |
| 187 | 492159.9196 | 7661487.299 | om |
| 193 | 492133.5337 | 7661782.781 | o |
| 199 | 492102.3307 | 7662084.346 | o |
| 247 | 492360.5752 | 7660497.17 | o |
| 265 | 492273.821 | 7661395.87 | om |
| 271 | 492241.3317 | 7661690.826 | om |
| 277 | 492210.8533 | 7661988.833 | om |
| 325 | 492466.3526 | 7660411.006 | m |
| 349 | 492350.7307 | 7661605.052 | om |
| 355 | 492318.0313 | 7661898.963 | om |
| 403 | 492579.6708 | 7660322.855 | om |
| 421 | 492489.8345 | 7661217.659 | o |
| 427 | 492459.9931 | 7661515.596 | o |
| 433 | 492429.7097 | 7661815.09 | o |
| 439 | 492400.5412 | 7662109.863 | om |
| 481 | 492685.3187 | 7660233.035 | om |
| 505 | 492570.4641 | 7661423.647 | om |
| 511 | 492538.6456 | 7661720.31 | om |
| 517 | 492510.545 | 7662016.268 | om |
| 577 | 492705.3132 | 7661035.606 | om |
| 589 | 492648.6405 | 7661627.855 | om |
| 595 | 492618.6039 | 7661928.998 | om |
| 667 | 492757.2465 | 7661545.107 | om |
| 679 | 492700.207 | 7662141.065 | om |
| 733 | 492924.5712 | 7660857.792 | o |
| 739 | 492896.5419 | 7661156.949 | om |
| 745 | 492866.4088 | 7661456.958 | om |
| 751 | 492838.1235 | 7661756.274 | om |
| 757 | 492808.1934 | 7662053.996 | o |
| 811 | 493035.5804 | 7660769.903 | o |
| 823 | 492977.5755 | 7661365.71 | om |
| 835 | 492917.5722 | 7661962.542 | om |
| 889 | 493144.4042 | 7660673.747 | o |
| 895 | 493112.3677 | 7660979.071 | om |
| 901 | 493083.4854 | 7661273.049 | om |
| 913 | 493025.7016 | 7661870.94 | o |
| 919 | 492997.2759 | 7662166.759 | o |
| 961 | 493283.5397 | 7660291.576 | o |
| 967 | 493251.6385 | 7660587.25 | om |
| 979 | 493194.3672 | 7661182.349 | o |
| 1045 | 493359.8784 | 7660495.105 | o |
| 1075 | 493213.6389 | 7661987.52 | om |
| 1123 | 493470.526 | 7660406.483 | om |
| 1159 | 493295.0253 | 7662195.316 | om |
| 11201 | 492845.6659 | 7660149.446 | o |
| 11203 | 492971.6613 | 7659907.651 | o |
| 11204 | 493012.2967 | 7659781.888 | m |
| 11205 | 493103.951 | 7659750.073 | o |
| 11207 | 493267.7171 | 7659911.392 | o |
| 11208 | 493180.9779 | 7660039.125 | o |
| 11209 | 492540.0029 | 7662347.483 | o |
| 11210 | 492576.8888 | 7662442.109 | o |
| 11211 | 492578.1461 | 7662627.43 | om |
| 11212 | 492584.1153 | 7662793.33 | om |
| 11213 | 492664.3919 | 7662852.22 | om |
| 11214 | 492751.5195 | 7662776.032 | o |
| 11215 | 492751.2244 | 7662647.351 | om |
| 11216 | 492750.4017 | 7662502.435 | om |
| 11220 | 493073.521 | 7661369.919 | m |
| 11221 | 492985.5579 | 7661243.017 | om |
| 11224 | 492593.5013 | 7661897.597 | om |
| 11225 | 492854.5453 | 7661994.276 | o |

**S2 Supplementary Figure 1.**
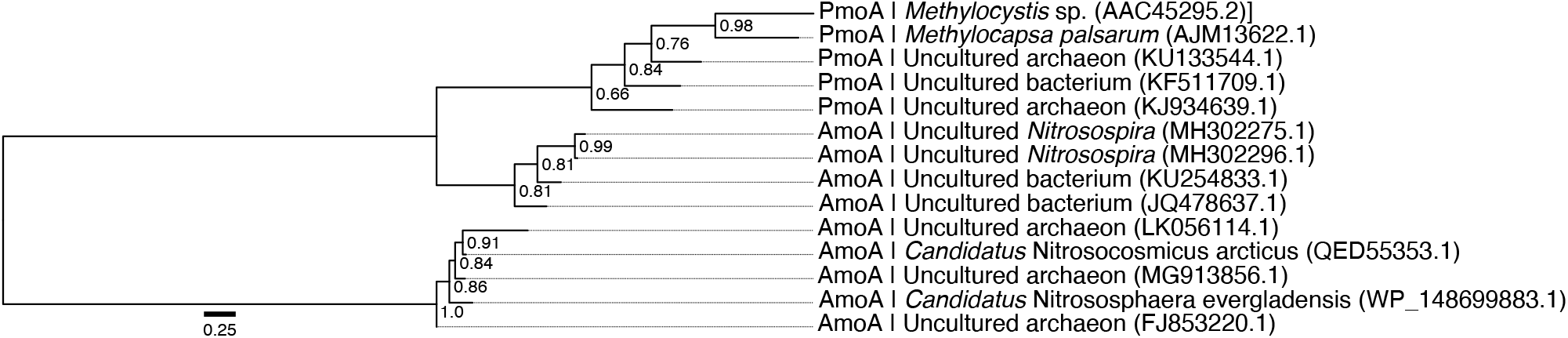
Maximum-likelihood tree of reference amino acid sequences used to discriminate between *amoA* and *pmoA* transcripts. Branch supports based on 1000 bootstraps are indicated.

**S3 Supplementary Figure 2.**
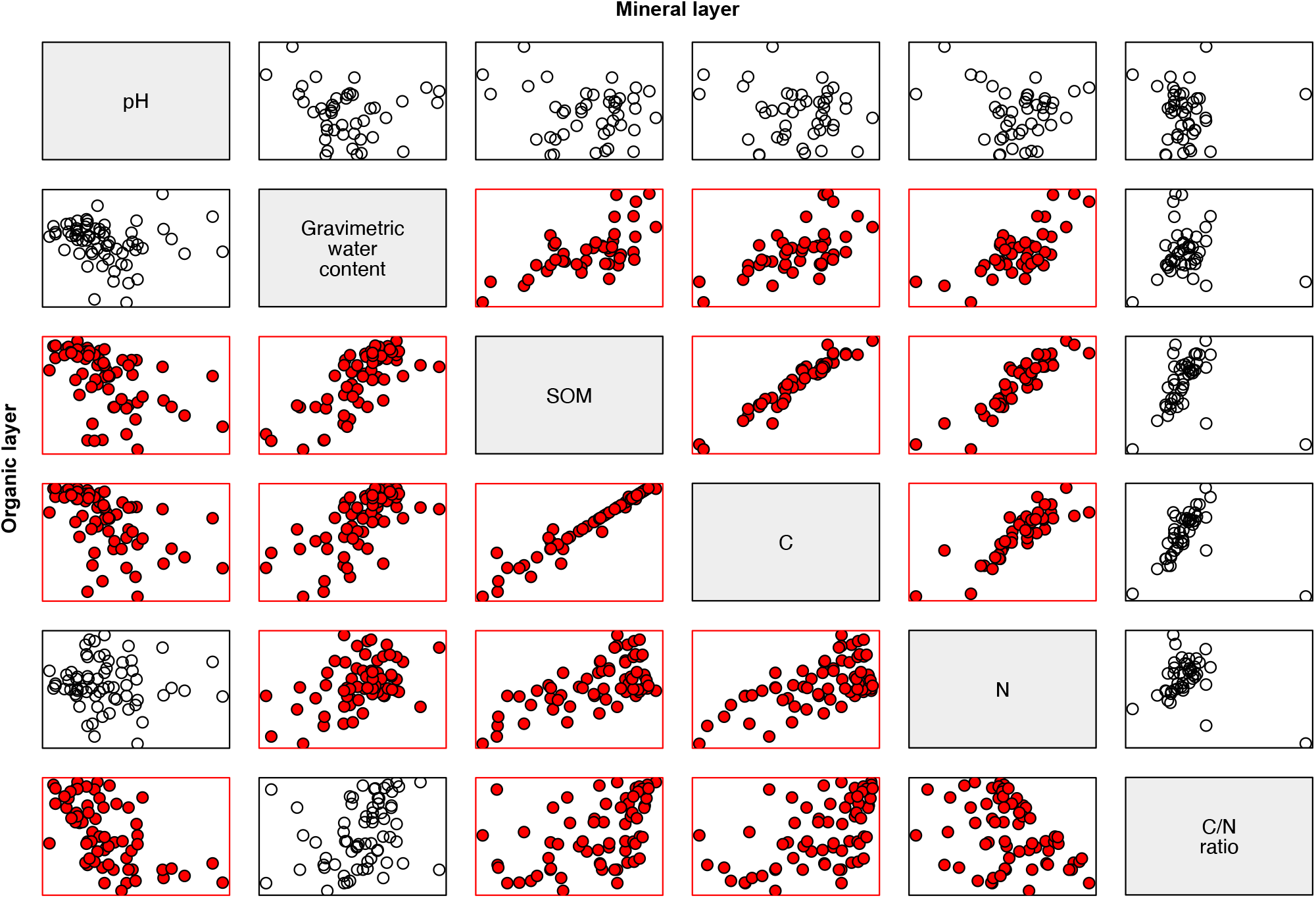
Pairwise correlation between soil physicochemical variables. Comparisons with Pearson correlation values (*r*) ≥ 0.5 or ≤ –0.5 are highlighted in red. SOM: soil organic matter; C: carbon; N: nitrogen.

**S4 Supplementary Table 2.**
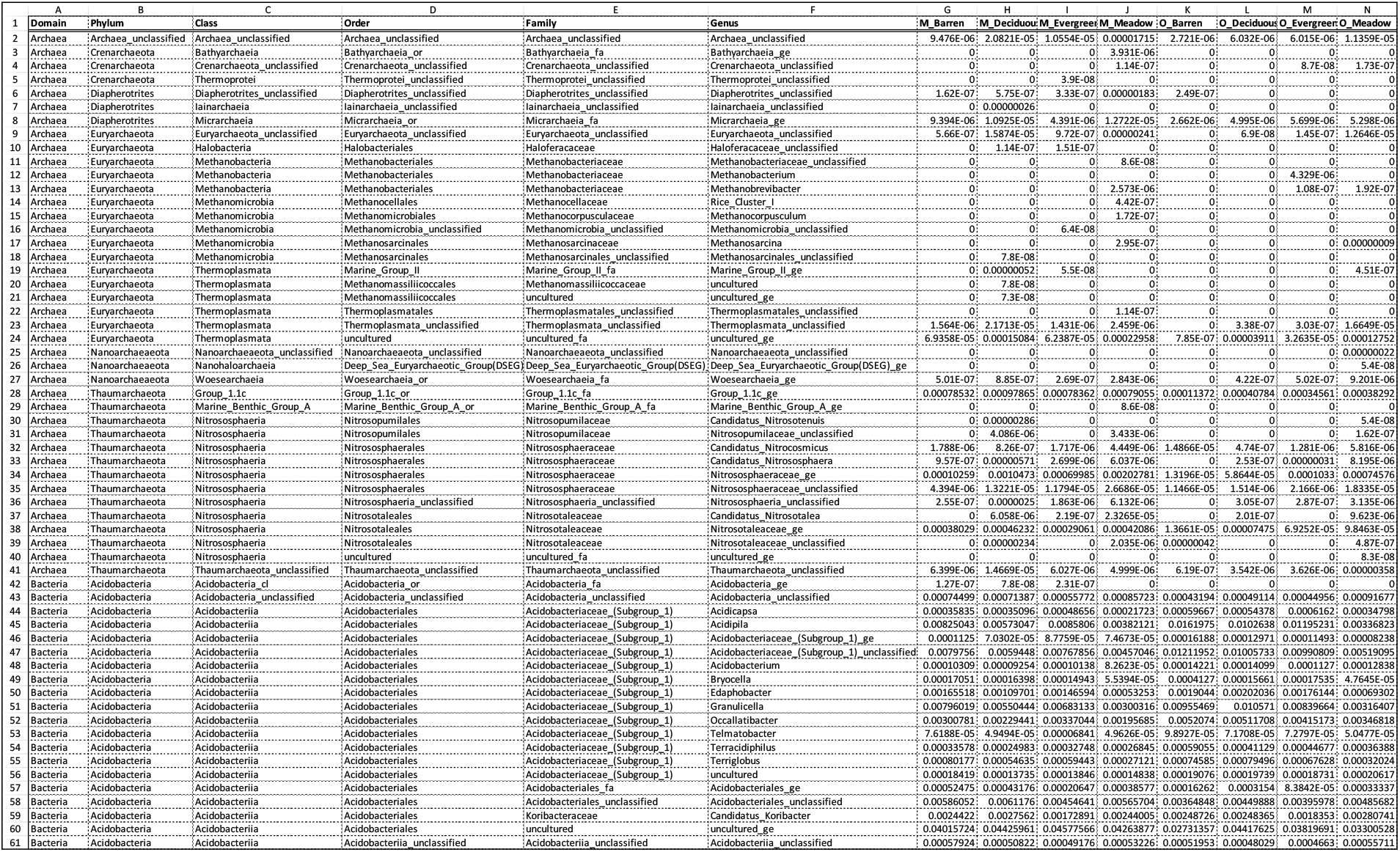
Relative abundances at the genus level in each vegetation type and soil layer.

**S5 Supplementary Figure 3.**
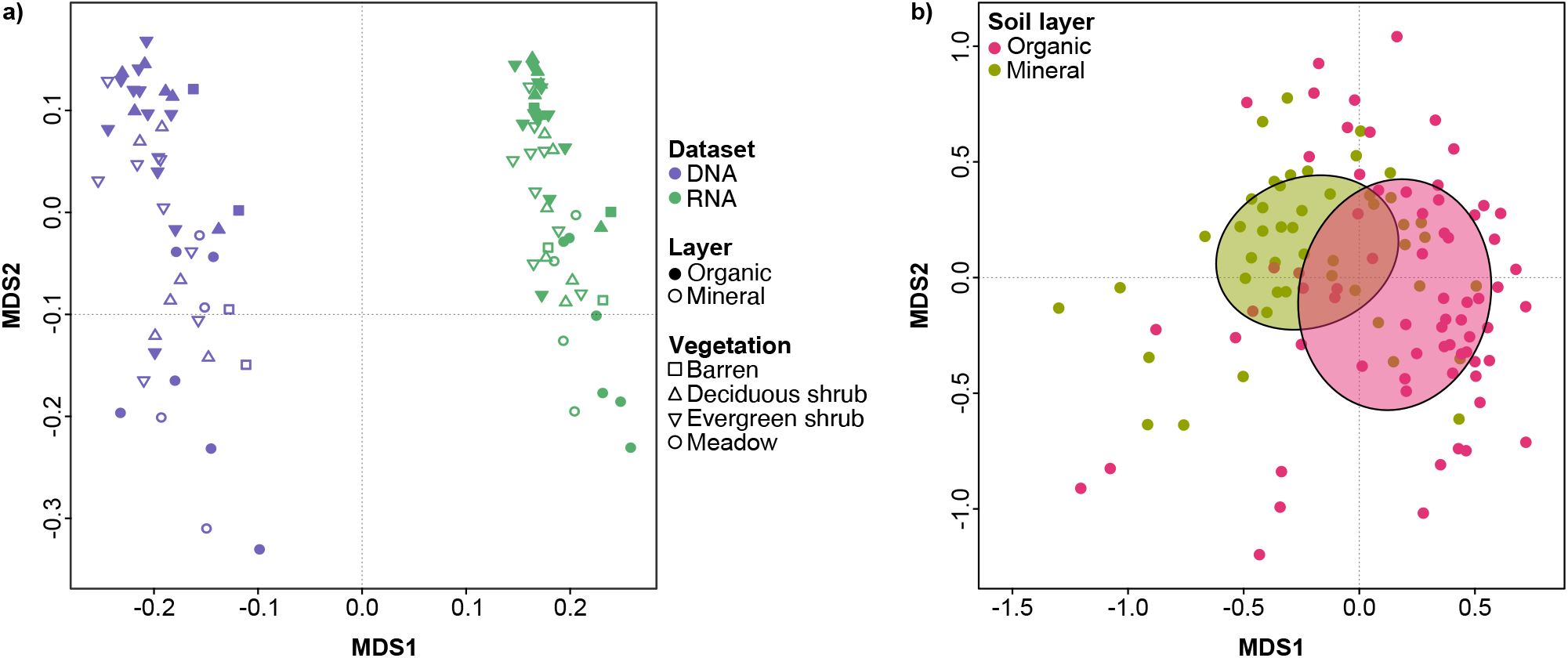
Principal coordinates analysis (PCoA) showing differences in genus-level taxonomic composition between **a)** metatranscriptomes (this study) and metagenomes (Pessi *et al*., 2021a) and **b)** soil layers in the metatranscriptomes. Ellipses in panel **b** represent 1.5 standard deviations from the group centroid.

**S6 Supplementary Table 3.**
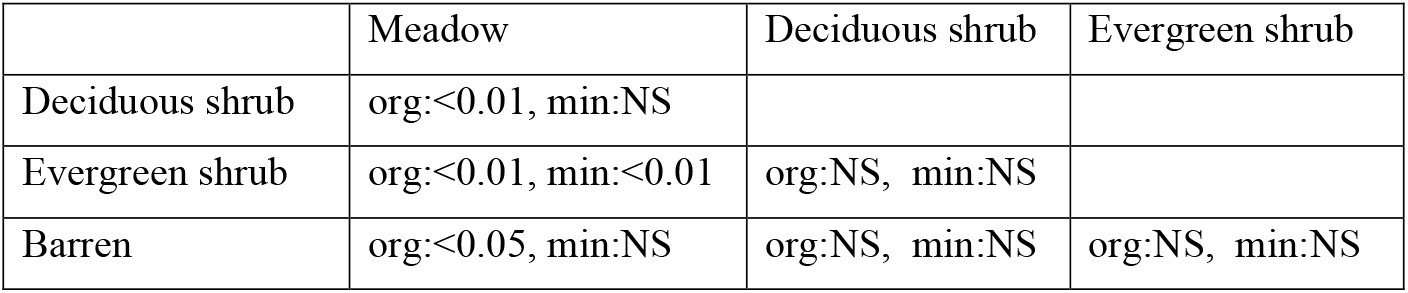
Differences in the active microbial communities between organic and mineral layers and the four different vegetation types based on pairwise PERMANOVA analysis. NS: not significant.

**S7 Supplementary Figure 4.**
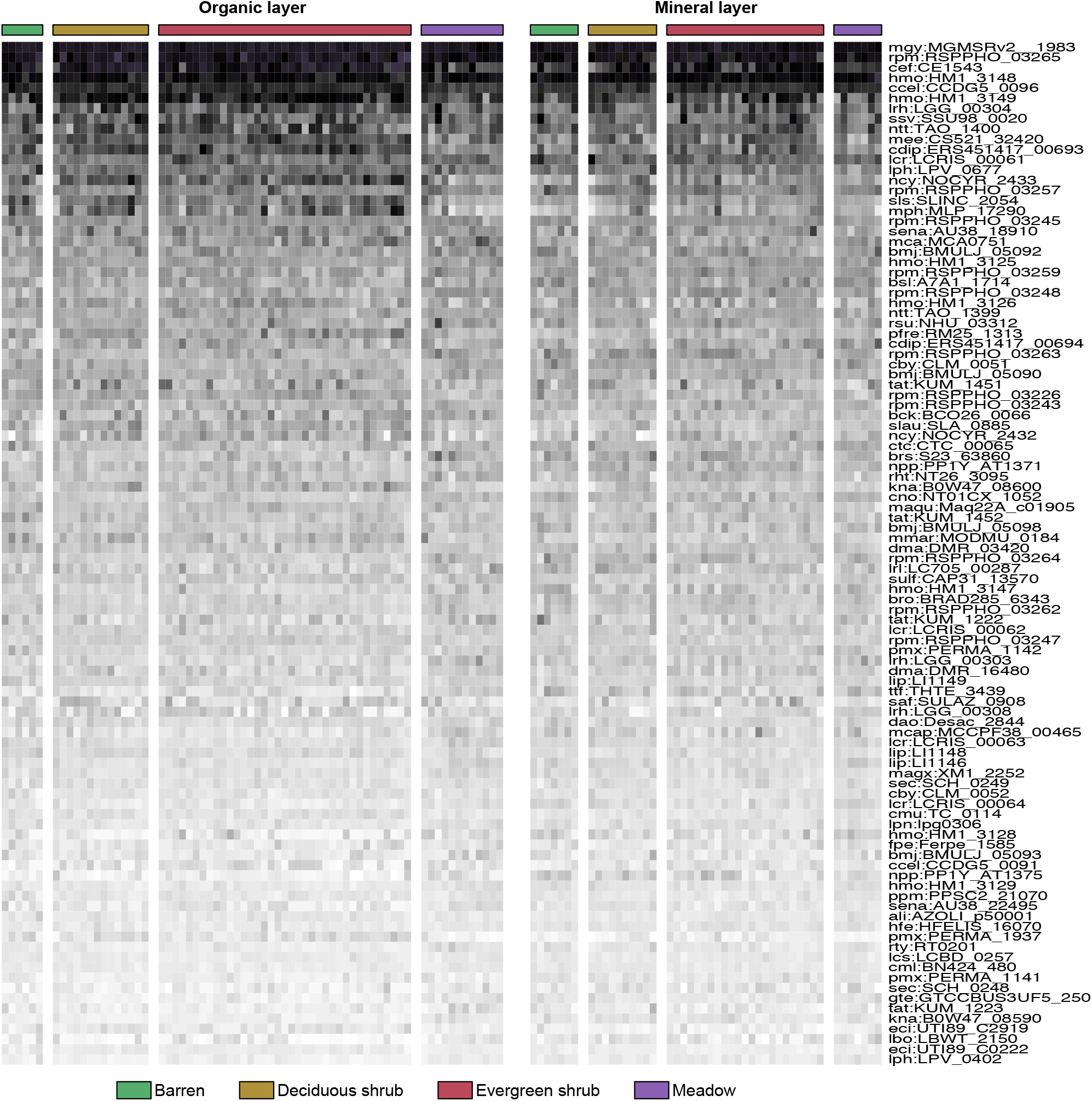
Heatmap showing the most abundant genes across all samples that did not match to any sequence present in the KEGG database. Abundances were square root-transformed to improve visualization.

**S8 Supplementary Figure 5.**
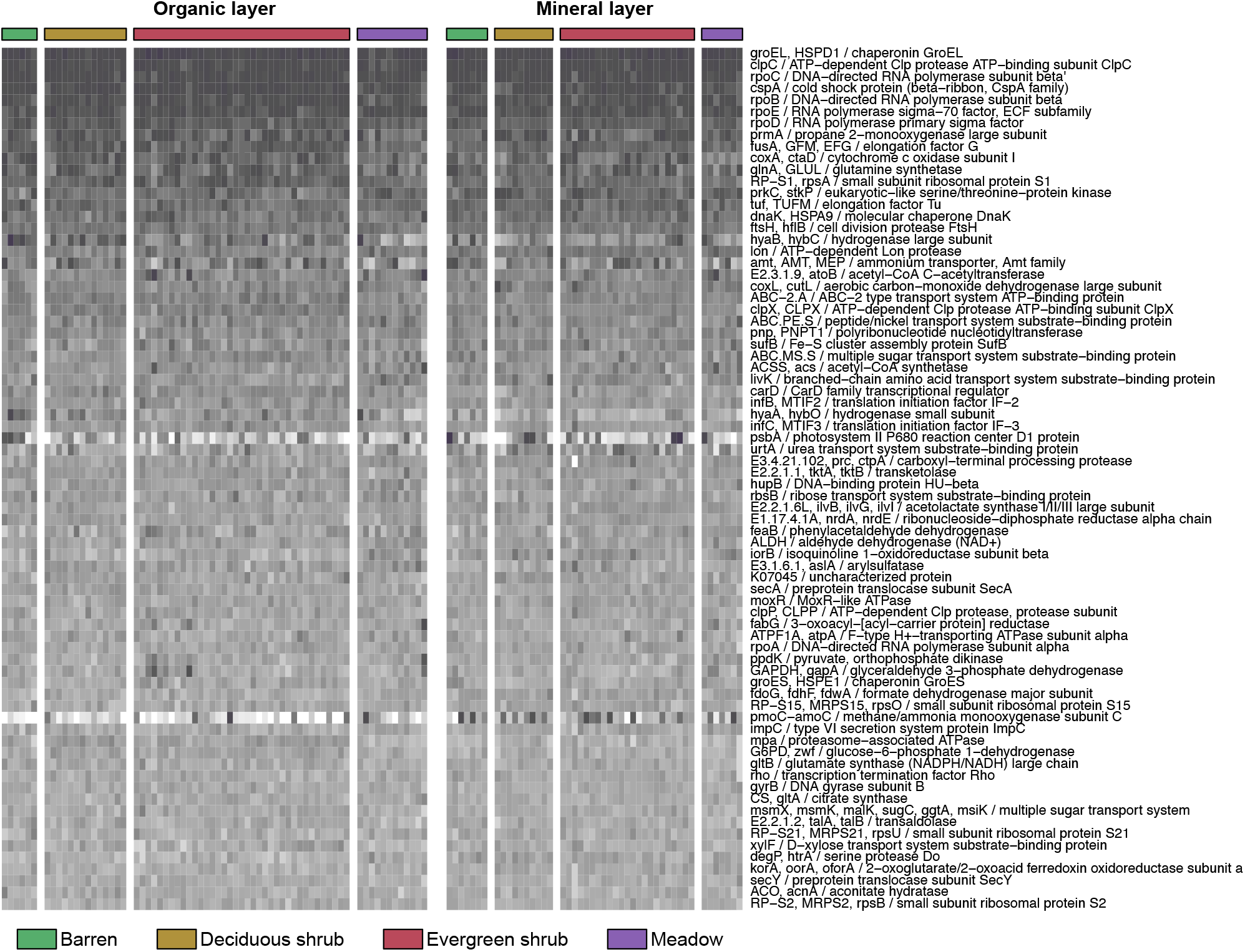
Heatmap showing the 100 most abundant genes that were mapped to KEGG pathways. Abundances were square root-transformed to improve visualization.

**S9 Supplementary Figure 6.**
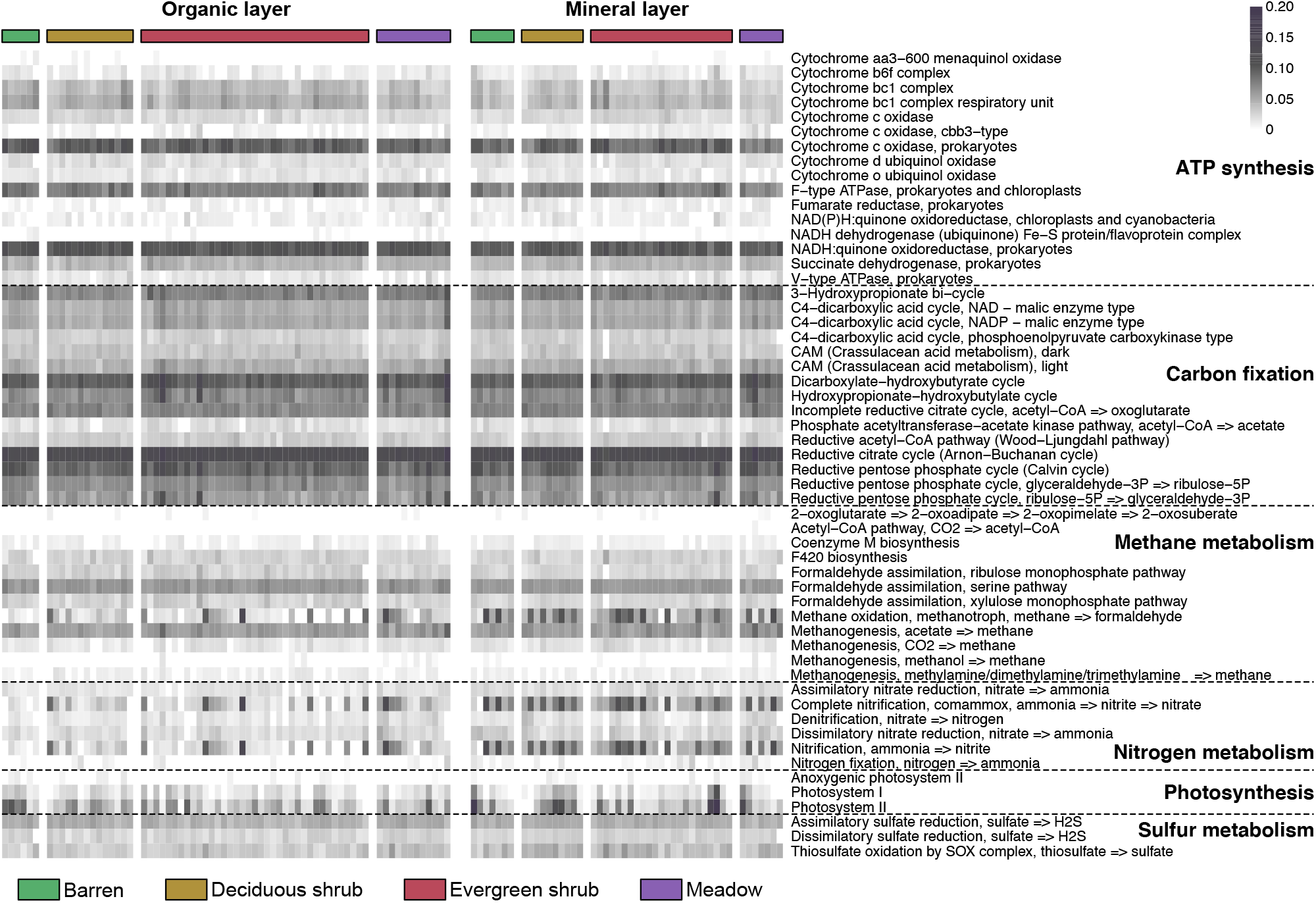
Heatmap showing the genes belonging to metabolism pathways in KEGG. Abundances were square root-transformed to improve visualization.

